# Engineering a highly durable adeno-associated virus receptor for analytical applications

**DOI:** 10.1101/2023.05.29.542019

**Authors:** Kouhei Yoshida, Yuji Tsunekawa, Kento Kurihara, Kazuya Watanabe, Yuriko Makino, Toru Tanaka, Teruhiko Ide, Takashi Okada

**Author notes:** Correspondence should be addressed to K.Y. and T.O.

## Abstract

Adeno-associated virus (AAV) is a major viral vector used in gene therapy. There are multiple AAV serotypes, and many engineered AAV serotypes with modified capsids altering tissue tropism are enthusiastically being developed. The universal AAV receptor (AAVR) is an essential receptor for multiple AAV infections. Since most AAV serotypes used in gene therapy infect cells via interaction with AAVR, the quantification of the vector-binding ability of AAV to AAVR could be an important quality check for therapeutic AAV vectors. To enable a steady evaluation of the AAV-AAVR interaction, we created an engineered AAVR through mutagenesis. Engineered AAVR showed high durability against acid while retaining its AAV-binding activity and an affinity chromatography column with engineered AAVR was also developed. This column enabled repeated binding and acid dissociation measurements of AAVR with various AAV serotypes. Our data showed that the binding affinities of AAV to AAVR were diverse among serotypes, providing insight into the relationship with the infection efficiency of AAV vectors. Thus, this affinity column can be used in process development for quality checks, quantitating capsid titers, and affinity purification of AAV vectors. Furthermore, this column may work for a useful tool of novel AAV vector capsid engineering.

## Introduction

The adeno-associated virus (AAV) contains a non-enveloped icosahedral capsid consisting of 60 subunits of 3 distinct viral proteins (VPs), approximately 25 nm in size, and a single-stranded DNA genome^1^. AAV is widely used as a vector for gene therapy owing to its low immunogenicity and cytotoxicity, persistent gene expression, and high efficiency in infecting various tissues^2–4^. The recent development of Zolgensma^®^, an AAV gene therapy containing the *SMN1* gene, has enabled the treatment of spinal muscular atrophy^5, 6^. Also, Roctavian and Hemgenix have been developed as the first AAV gene therapy containing the gene for factor VIII and IX to treat hemophilia A and B^7^. In addition, AAV vectors carrying the gene for microdystrophin are being developed for Duchenne muscular dystrophy treatment^8–11^. Therefore, there are high expectations for the development of AAV as a breakthrough treatment for intractable hereditary diseases.

AAVs utilize various receptors to infect cells^12^. Among these, the receptor expressed by the *KIAA0319L* gene, identified as a dyslexia-associated protein, mainly contributes to binding and infection in various AAVs and is known as the universal AAV receptor (AAVR)^13, 14^. AAVR-knockout cells acquired resistance to most AAV infections, indicating the importance of evaluating the binding of AAV and AAVR to understand the performance of AAV in gene therapy^14^. AAVR is also being investigated for its application as an affinity ligand, including its use as an AAV carrier^15^.

In this study, we aimed to expand the availability of AAVR for the analytical application of AAV *in vitro* by improving the durability of AAVR. We created an AAVR mutant using an evolutionary engineering method, enabling the repeated evaluation of the binding of AAVs while retaining their function. We developed an AAV affinity resin with an AAVR mutant and assessed its performance using a column packed with it. This column enables several AAV assessments using an HPLC system.

## Results

### Creation and evaluation of durable AAVR mutant

AAVR is a membrane protein with five extracellular polycystic kidney disease (PKD) domains (PKD1–5), of which PKD1 and PKD2 are highly involved in AAV binding^16^. Therefore, we selected PKD1 and PKD2 from AAVR as the AAV affinity ligands. The wild-type (WT)-AAVR was added to an SPR sensor chip immobilized with AAV2 and its functional activity was confirmed by the binding response (Fig. 1A). Enzyme-linked immunosorbent assay (ELISA) was used to measure the durability of WT-AAVR after soaking it in acid, which showed a decrease in AAV-binding activity with soaking time (Fig. 1B). Therefore, it was deemed necessary to improve the durability of WT-AAVR in acids by introducing mutations.

**Fig. 1.**
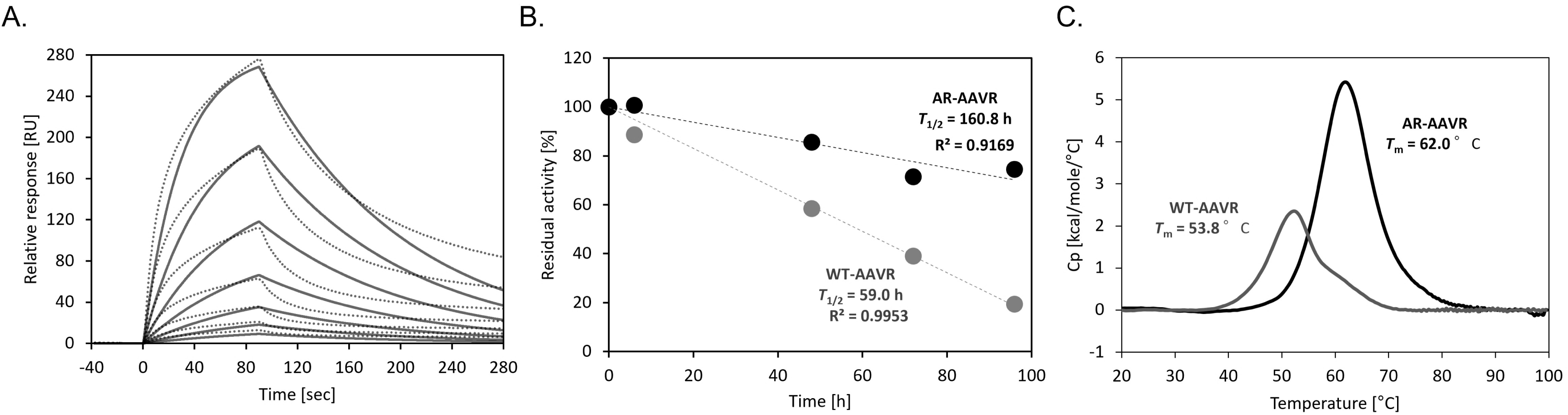
(A) Binding response of the interaction between AAV2 and WT-AAVR in SPR. (B) Plot of residual activity of acid-soaked WT-AAVR (gray) and AR-AAVR (black) against AAV2 over time. The black and gray dotted lines represent linear trendlines calculated from each plot. The R-squared value of the trendlines is shown in the graph. *T*_1/2_ indicates the time at which the residual activity reached 50%. (C) Melting curves of WT-AAVR (gray) and AR-AAVR (black) and calculated *T*_m_ values in DSC.

Error-prone PCR was performed using the gene sequence encoding WT-AAVR to generate a cDNA library containing random mutations^17^. This library was transformed into *Escherichia coli*, and the expressed mutants were harvested from the culture supernatant. After acid treatment (0.1 M glycine-HCl; pH 3.0) of the mutants, their interaction with AAVs was evaluated through ELISA. While the WT-AAVR lost its AAV-binding activity, several clones maintained their AAV-binding activity (data not shown). Consequently, an acid-resistant (AR)-AAVR was created by introducing 10 mutations that conferred acid durability into WT-AAVR. AR-AAVR showed a 2.5-fold longer half-life in acid than did WT-AAVR (Fig. 1B). In addition, during differential scanning calorimetry (DSC) assessments, AR-AAVR exhibited 8°C higher *T*_m_ than did WT-AAVR, indicating that these introduced mutations contributed to the rigidity and stability of AAVR and also improved its durability in acids (Fig. 1C).

However, the introduction of random evolutionary mutations into proteins may result in the loss of their intrinsic binding properties. AAVR has been reported to exhibit two modes of interaction while binding to different AAV serotypes^12, 18–20^. In most AAV serotypes, such as AAV1, AAV2, and AAV9, AAVR binds along the 3-fold symmetry axis of AAVs. However, AAV5 is phylogenetically distant from other AAV serotypes and has low amino acid sequence homology; therefore, unlike other AAVs, AAVR binds to AAV5 along a 5-fold symmetry axis. PKD1 and PKD2 of AAVR can recognize and interact with AAV through these two distinct binding modes. To confirm this ability of AAVR, its capacity to bind to AAV2 and AAV5 was measured using surface plasmon resonance (SPR). AR-AAVR retained its binding ability, indicating that stabilization mutations did not affect AAV binding (Fig. S1).

### Durability comparison of AAVR affinity column

Affinity resins were prepared by anchoring WT-AAVR or AR-AAVR to a non-porous resin and packing them into SUS columns (Figs. 2A and 2B). These columns were connected to an HPLC instrument, and purified AAV2 was injected into the columns. The captured AAV2 was eluted at approximately 30 min for WT-AAVR and 40 min for AR-AAVR through the mobile phase B gradient and detected by fluorescence (Figs. 2C and 2D). After repeating this measurement 100 times, the chromatograms obtained using WT-AAVR showed that the peak derived from the eluted AAV2 gradually became smaller and broader (Fig. 2C). In addition, fluorescence increased immediately after injection, representing the unbound AAV2 on the column, suggesting that the performance of WT-AAVR was insufficient and decreased with repeated use. In contrast, the column using AR-AAVR (hereafter, referred to as “AVR-NPR column”) maintained the sharpness and size of the AAV2 elution peak even after the 100^th^ assessment (Fig. 2D). In addition, the very little flow-through of AAV2 immediately after injection indicated a strong binding ability to AAV and the high durability of the column.

**Fig. 2.**
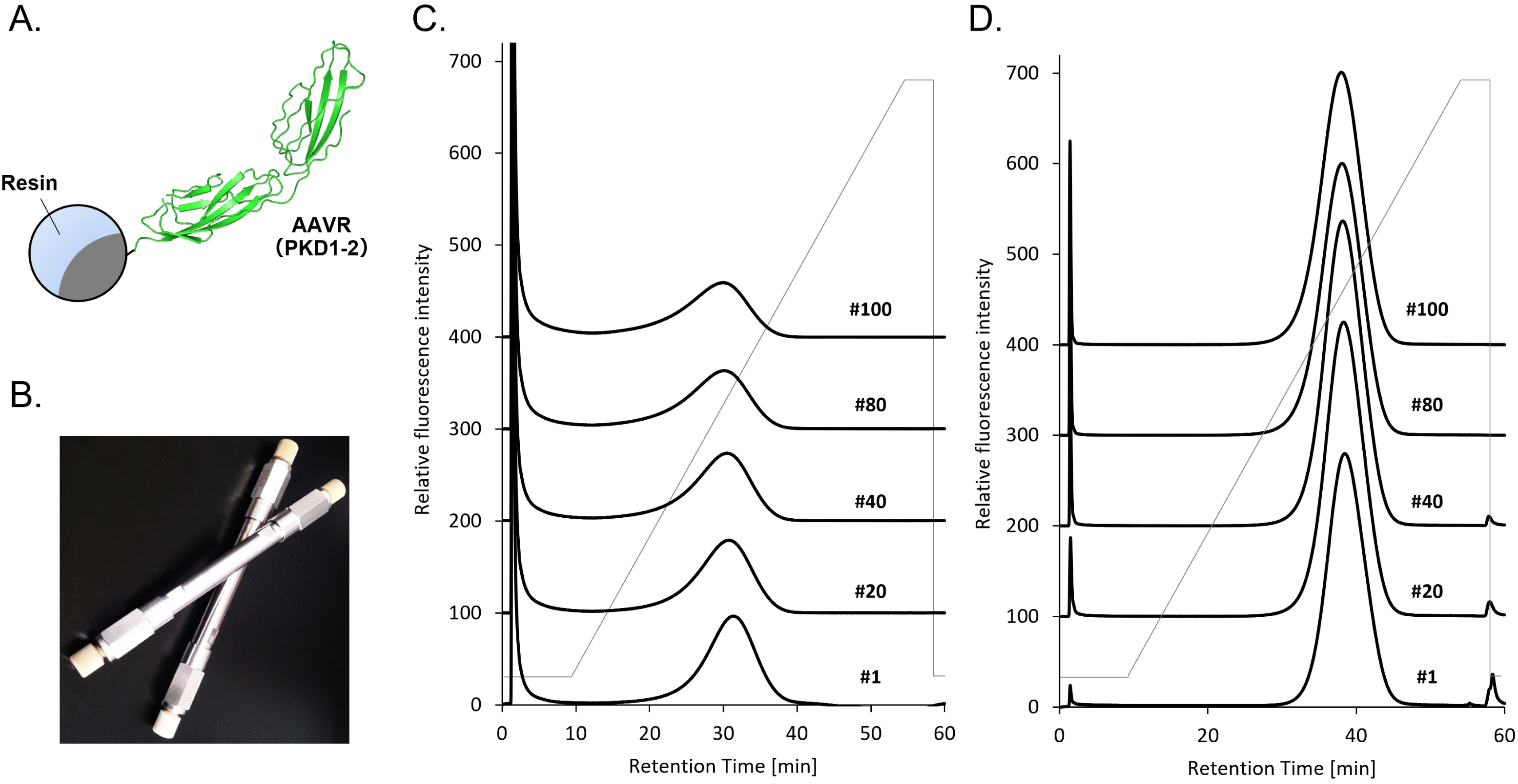
(A) Schematic diagram of the affinity resin. The ribbon diagram of the ligand was generated by AlphaFold2. (B) The outside appearance of the SUS column. (C, D) Chromatograms when purified AAV2 was subjected to the (C) WT-AAVR or (D) AR-AAVR affinity column. The numbers on the right side of each chromatogram indicate the number of repeated trials. The black lines indicate the relative fluorescence intensity at each time. The gray polygonal line represents the ratio of mobile phase A (15 mM sodium acetate, 10 mM glycine, and 50 mM CaCl_2_ (pH 4.5)) and B (15 mM sodium acetate, 10 mM glycine, and 50 mM CaCl_2_ (pH 2.2)), which here the ratio of B increases linearly from 0% to 100% between 10 min and 55 min after the start of the measurement.

### Analysis of various AAVs with AVR-NPR column

AAV1, AAV2, AAV4, AAV5, AAV8, and AAV9, which have different tissue tropisms, were expressed using a suspension cell expression system and purified by affinity chromatography.

AAV1, AAV2, AAV5, AAV8, and AAV9 were captured on an AVR-NPR column and eluted using a pH gradient (Fig. 3A). AAV4 was previously shown not to bind to AAVR^21^ and was not consistently captured in this column (data not shown). The difference in the retention times of the elution peaks for each AAV serotype could be due to the difference in the affinity of the interaction with AAVR, suggesting the possibility of its application in AAV identification and capsid modification.

**Fig. 3.**
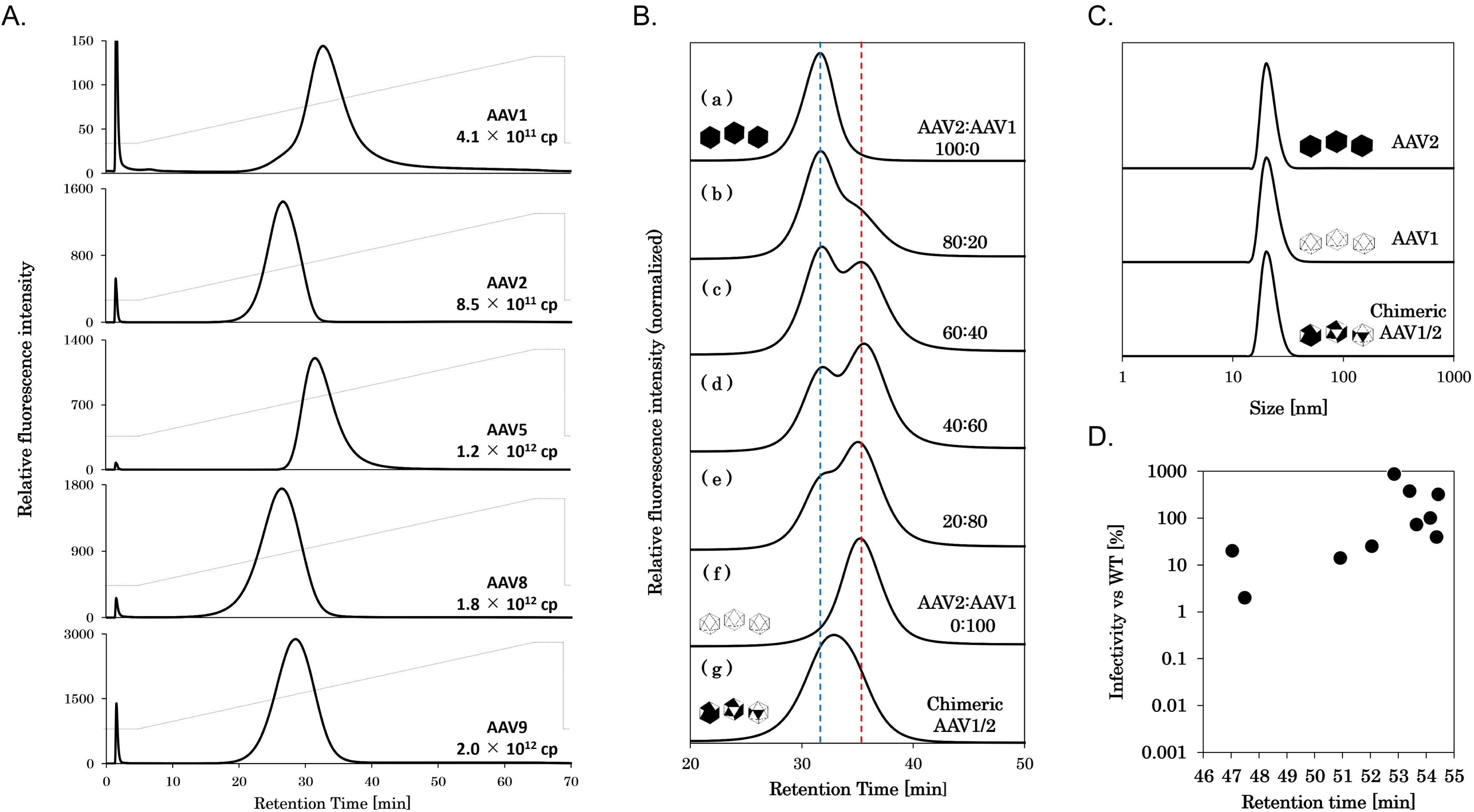
(A) Chromatograms when purified AAVs were subjected to the AVR-NPR column. The AAV serotypes and injection amounts are indicated on each chromatogram’s right side. The black lines indicate the relative fluorescence intensity, and the gray polygonal line represents the ratio of mobile phase A (15 mM sodium acetate, 10 mM glycine, and 50 mM CaCl_2_ (pH 4.5)) and B (15 mM sodium acetate, 10 mM glycine, and 50 mM CaCl_2_ (pH 2.2)), which the ratio of B increases linearly from 0% to 100% between 5 min and 65 min after the start of the measurement. (B) Chromatograms when (a-f) a mixture of AAV1 and AAV2 in different ratios and (g) chimeric AAV1/2 were subjected to the AVR-NPR column. The black lines indicate the relative fluorescence intensity. The vertical red and blue dashed lines represent the top elution peak positions of AAV1 and AAV2, respectively. (C) Particle sizes of AAV1, AAV2, and AAV1/2 in DLS. (D) Plots of the retention time against the transduction efficiency of each AAV2 mutant from Table 1 when analyzed using the AVR-NPR column.

**Table 1.**
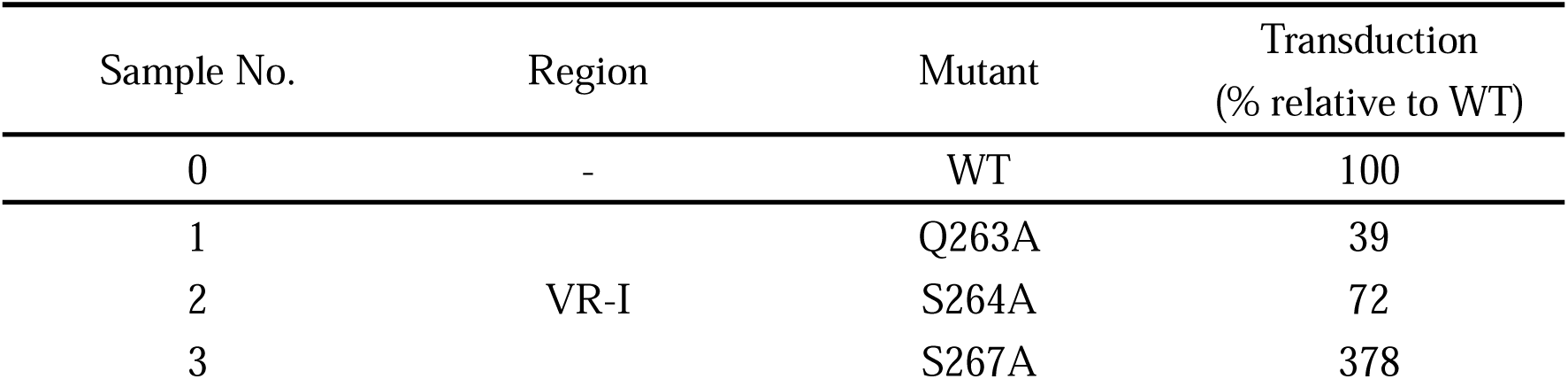
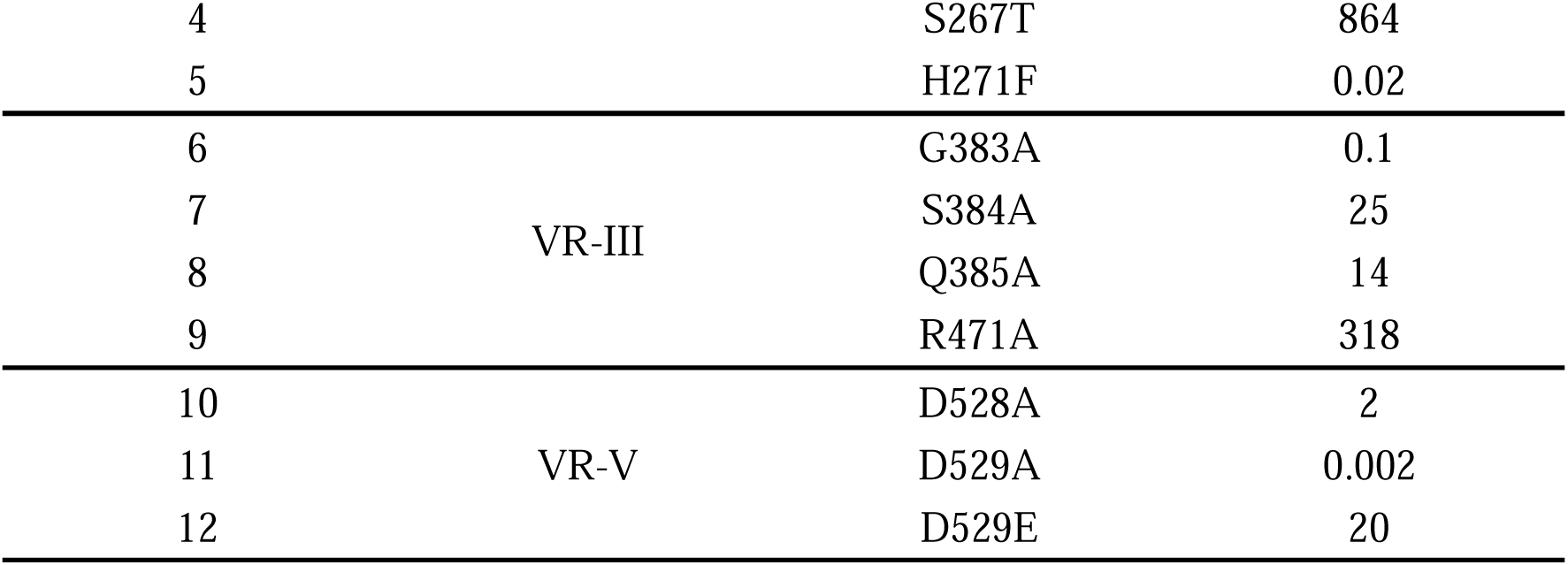
List of AAV2 mutations. VR locations of each mutation and the transduction efficiency of each mutant relative to the WT are indicated.

To test this hypothesis, we conducted chimeric and mutant AAV analyses using an AVR-NPR column. When mixtures of AAV1 and AAV2 in different ratios were subjected to column analysis, the bimodality of the detection peaks varied with the proportion of each AAV serotype (Fig. 3B). By mixing plasmids carrying the Cap genes of AAV1 and AAV2 and expressing AAV, chimeric AAV (AAV1/2) can be produced, in which the VPs from each AAV are mixed to form AAV particles^22^. When AAV1/2 was analyzed using the AVR-NPR column, a broad elution peak was observed, in contrast to the peak for the AAV1 and AAV2 mixture, suggesting that the surface structure of AAV1/2 was heterogeneous depending on the proportion of the serotypes, resulting in the production of chimeric AAVs with various binding affinities to AAVR (Fig. 3B). Dynamic light scattering (DLS) of AAV1, AAV2, and AAV1/2 revealed similar particle sizes and shapes, indicating that AAV heterogeneity cannot be distinguished (Fig. 3C). Therefore, the AVR-NPR column is unique in its ability to distinguish slight differences in the capsid surface structure of AAV serotypes, based on the affinity of the AAV-AAVR interaction.

Moreover, point mutations that alter AAV2 infectivity have previously been reported^23^, and we extracted 12 AAV2 mutations that appear to be involved in the interaction with AAVR (Table 1). These AAV2 mutants were subjected to AVR-NPR column analysis, which showed a correlation between the retention time of the elution peak and the infectivity of the mutants (Fig. 3D). Mutants with short retention times showed a reduced affinity for AAVR, which resulted in reduced cell infectivity. Three AAV2 mutants (H271F, G383A, and D529A) whose infectivity was reduced to less than 0.1% compared with that of the WT did not show elution peaks, suggesting that these mutants were not captured by the AVR-NPR column. In comparison, all AAV2 mutants were captured using an affinity column with a non-AAVR protein as a ligand, and no such correlation was observed (Fig. S2). Therefore, the AVR-NPR column can be an effective tool for evaluating the infectivity of newly created AAVs.

### AAV quantification using the AVR-NPR column

Various amounts of purified AAV8 were analyzed using the AVR-NPR column (Fig. 4A) and the elution peak area of each measurement was plotted against the actual particle amount. These were proportional with high linearity in the range of 10^9^–10^12^ cp of the AAV8 quantity (Fig. 4B).

**Fig. 4.**
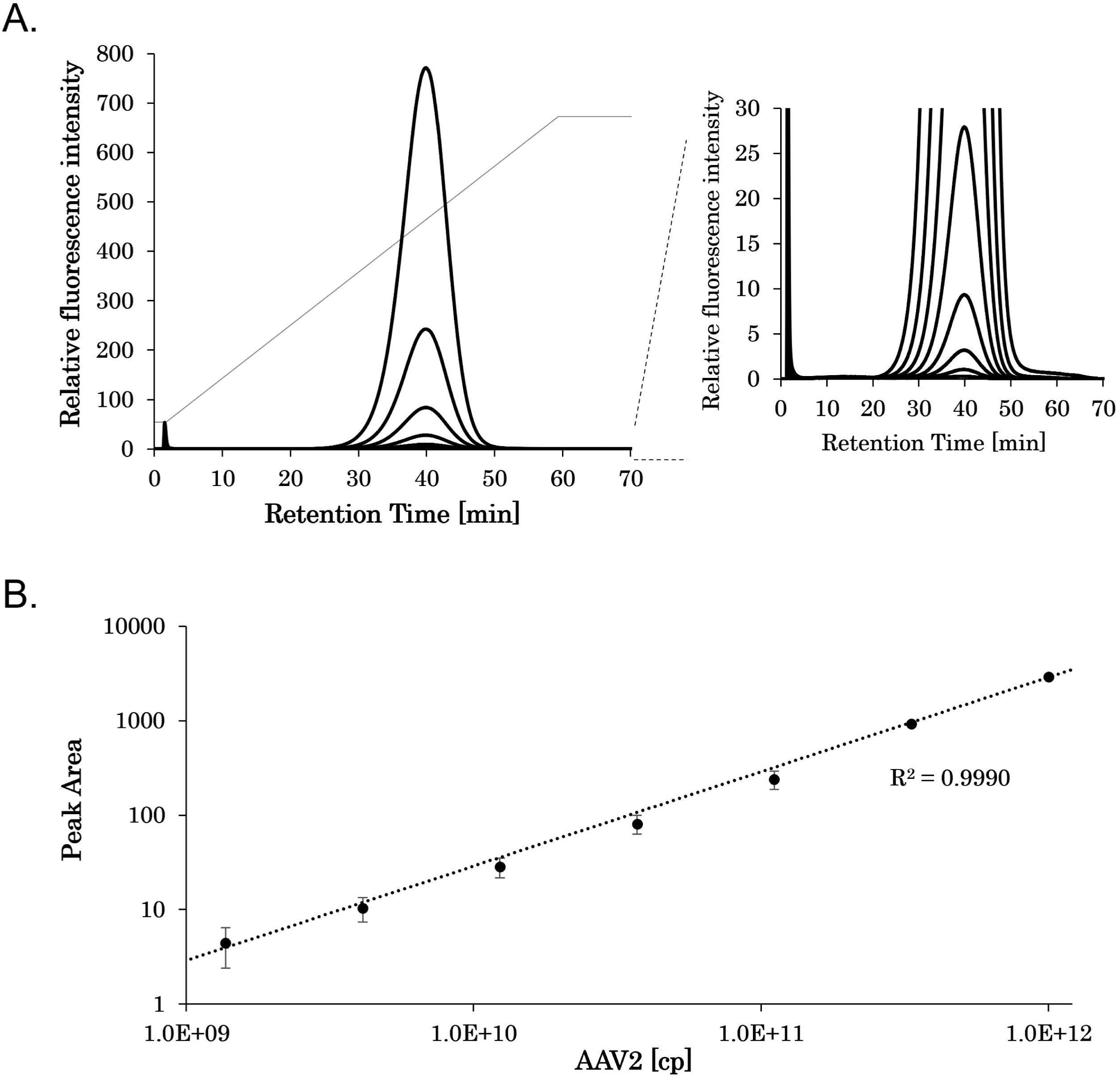
Quantification of AAV8 particle amount using the AVR-NPR column. (A) Chromatograms when purified AAV8 particles were injected at 1.0 × 10^12^ cp, 3.3 × 10^11^ cp, 1.1 × 10^11^ cp, 3.7 × 10^10^ cp, 1.2 × 10^10^ cp, 4.1 × 10^9^ cp, and 1.4 × 10^9^ cp. The black lines indicate the relative fluorescence intensity, and the gray polygonal line represents the ratio of mobile phase A (15 mM sodium acetate, 10 mM glycine, and 50 mM CaCl_2_ (pH 4.5)) and B (15 mM sodium acetate, 10 mM glycine, and 50 mM CaCl_2_ (pH 2.2)), which the ratio of B increases linearly from 0% to 100% between 3 min and 63 min after the start of the measurement. A zoom view of the chromatograms is shown in the right window. (B) Plots of the peak areas of the chromatogram from (A) against the AAV8 injection amount. The assessments were performed in triplicate, and the error bars indicate the standard deviation. The black dotted line indicates the linear trendline calculated from the plots. The R-squared value of the trendline is shown in the graph.

When the cell-lysed supernatants were subjected to AVR-NPR column analysis, medium-and cell-derived impurities passed through the column immediately after injection, and the fluorescence was saturated over the detectable limit. In the AAV8-expressing supernatant, an AAV8-derived elution peak was observed at approximately 75 min, whereas for the untreated supernatant, an elution peak was not observed (Fig. S3). Thus, the AVR-NPR column specifically captured and detected AAV expressed in the culture medium. By plotting the peak area of the detected AAV on the calibration curve described above, the capsid titer of AAV in the culture medium can be quantified, which can be used to monitor AAV expression.

To test the effect of host cell protein contamination during the AAV capsid titer monitoring with AVR-NPR column, purified AAV8 was mixed in cell-cultured or cell-lysed supernatants, and the concentration was quantified using the AVR-NPR column and ELISA for comparison. The AVR-NPR column produced a constant elution peak, irrespective of the degree of purification (Fig. 5A), allowing concentration quantification with high reproducibility. In contrast, ELISA tended to underestimate the concentration of AAV8 in the crude state, implying that impurities inhibited its interaction with AAV (Fig. 5B).

**Fig. 5.**
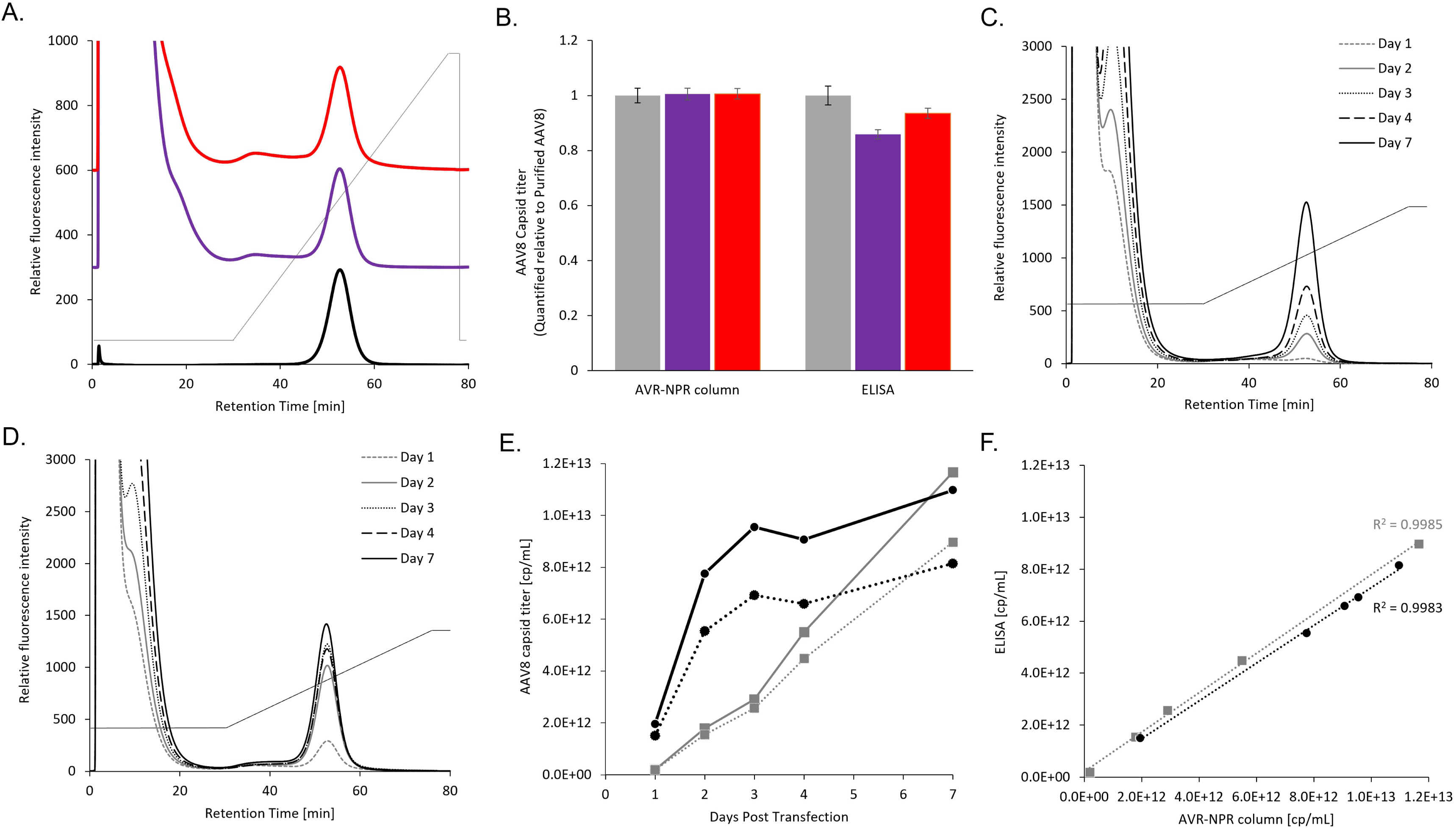
(A) Chromatograms of the purified AAV8 (black), AAV8-mixed cell-cultured supernatant (purple), and AAV8-mixed cell-lysed supernatant (red). The gray polygonal line represents the ratio of mobile phase A (15 mM sodium acetate, 10 mM glycine, and 50 mM CaCl_2_ (pH 4.5)) and B (15 mM sodium acetate, 10 mM glycine, and 50 mM CaCl_2_ (pH 2.2)), which here the ratio of B increases linearly from 0% to 100% between 30 and 75 min after the start of the measurement. (B) Quantification of AAV8 using the AVR-NPR column and ELISA. The bars show the purified AAV8 (gray), AAV8-mixed cell-cultured supernatant (purple), and AAV8-mixed cell-lysed supernatant (red). The capsid titers quantified with each purified AAV8 are shown as 1 and the other capsid titers as relative values. The analyses were performed in triplicate, and the error bars indicate the standard deviation. (C-F) Expression monitoring of AAV8. Chromatograms of (C) cell-cultured and (D) cell-lysed supernatants expressing AAV8 at days 1, 2, 3, 4, and 7 after the transfection. The gray polygonal line represents the ratio of mobile phase A and B, which here the ratio of B increases linearly from 0% to 100% between 30 and 75 min after the start of the measurement. (E) Quantification of the AAV8 capsid titers for each sample in (C) and (D). The results of cell-cultured supernatants quantified by the AVR-NPR column and ELISA are plotted as gray squares with solid and dotted lines, respectively. In contrast, the results of cell-lysed supernatants quantified by the AVR-NPR column and ELISA are plotted as black circles with solid and dotted lines, respectively. (F) Correlation functions between the AAV8 capsid titers quantified by the AVR-NPR column and ELISA. Gray squares and black circles were plotted based on the quantification results of cell-cultured and cell-lysed supernatant in (E), respectively. Each dotted line indicates the linear trendline calculated from the plots. R-squared values of the trendlines are shown in the graph.

### Monitoring AAV expression level in the culture

After transfection with AAV8, the culture medium was collected. The cell-cultured and cell-lysed supernatants were prepared via centrifugation and filtration, with or without the addition of a surfactant. In both cases, the AAV8-derived elution peak increased with the number of days post-transfection (Figs. 5C and 5D). The peak in the cell-lysed supernatant became saturated after day 3, whereas the peak in the cell-cultured supernatant gradually increased until day 7, suggesting that the ratio of extracellularly secreted AAV8 increased with an increase in the culture time after transfection. The transition of capsid titers of AAV8 in each supernatant, as calculated from the calibration curve, was similar to that measured using ELISA (Figs. 5E and 5F).

## Discussions

Although AAV utilizes many kinds of co-receptors to infect the cells, AAVR is one of the most important targets to evaluate AAV infectivity, so understanding AAV-AAVR interaction property would be an insight into better AAV vector development. In this study, we generated AAVR modified through mutagenesis, which showed improved durability without the loss of its AAV-binding function (Figs. 1, 2 and S1). The melting curve peak of AR-AAVR in the DSC measurement was larger than that of WT-AAVR (Fig. 1C), resulting in increased intramolecular interactions by introducing the mutations. The durability of the AR-AAVR enabled its use in analytical applications as an AAV affinity ligand, allowing the affinity evaluation of various AAVs with chromatography as an AVR-NPR column (Figs. 3A, 3B and 3D). The result that AAV-AAVR interaction affinity varied by serotype implied the possibility of a relationship between the strength of the interaction affinity and the pharmacokinetics of the AAV vector, such as diffusibility in the body. However, the relationship between affinity to the AVR-NPR column and cell infectivity for each AAV serotype remains to be elucidated. Although DLS is a powerful analytical tool in measuring AAV particle size distributions^24^, it cannot distinguish the surface structure of modified AAV by capsid engineering (Fig. 3C). Meanwhile, the AVR-NPR column could be used for a structural validation of newly created AAVs.

Also, the AVR-NPR column was available as a novel tool for AAV capsid quantification (Figs. 4 and 5). Our results showed that wide-ranged AAV capsid titer was easily assessed just by passing the AVR-NPR column through. This quantification range depended mainly on the sensitivity of the detector; for example, a more sensitive fluorescent detector can be used to quantify samples with lower capsid titers, or a UV detector instead of fluorescence can be used to quantify larger amounts of AAV. In addition, since the injection volume can be adjusted automatically in HPLC, the amount of AAV can be increased or decreased according to the capsid titer of the AAV sample to be measured, eliminating the concentration or dilution steps during pretreatment. Also, the ability of the AVR-NPR column to remove impurities through affinity chromatography would be advantageous for improving reproducibility (Fig. S3), and this could be an alternative AAV capsid quantification system to ELISA. Therefore, the AVR-NPR column can be used to easily quantify capsid titers of various AAV serotypes in the culture, which may provide an investigation into the culture conditions to harvest AAV vectors efficiently.

The AVR-NPR column would enable the accelerated development of gene therapy using AAVs. For example, capsid-engineered AAVs with various affinities to AAVR would be easily screened using the AVR-NPR column. Moreover, this column could be applied for affinity purification of multiple AAV serotypes. Since AAV vectors purified by interaction with AAVR ensure biological activity and improve their homogeneity, the AVR-NPR column will provide higher-quality AAV vector manufacturing platform.

## Material and methods

### Cloning, expression, and purification of AAVR

The amino acid sequences of the WT-AAVR and AR-AAVR are shown in Fig. S4. The DNAs encoding WT-AAVR and AR-AAVR were expressed in the vector pET-26b(+) (Merck), displaying a PelB signal peptide at the N-terminus and a hexa-histidine tag and cysteine at the C-terminus. *Escherichia coli* strain BL21 (DE3) cells (Novagen) were transformed using the expression vector and grown at 37°C in a 2× YT broth. When the optical density of the culture medium at 600 nm reached 0.6, isopropyl β-D-1-thiogalactopyranoside was added (0.1 mM).

The culture broth was harvested after 16 h at 20°C. The cells were harvested via centrifugation at 8,000 × g for 10 min, and the precipitate was resuspended in a solution containing 0.15 M NaCl, 20 mM imidazole, and 20 mM Tris-HCl (pH 7.4) (buffer A). The harvested cells were subsequently lysed via sonication for 15 min using a cell disruptor (Kubota Corporation, Tokyo, Japan). A precipitate containing soluble intracellular components was obtained through centrifugation at 12,000 × g for 20 min. The soluble fraction was collected and fed into Ni-NTA column (Cytiva, Tokyo, Japan) equilibrated with buffer A. The proteins were eluted with 500 mM imidazole contained in buffer A and then dialyzed with a solution containing 0.15 M NaCl and 20 mM Tris-HCl (pH 7.4). The concentrations of the purified proteins were measured using NanoDrop spectrophotometer (Thermo Scientific).

### Acid-resistance evaluation using ELISA

Purified proteins (0.01 mg/mL) were incubated in a solution of 0.1 M glycine (pH 3.0) at 30°C for 0, 6, 48, 72, and 96 h, followed by neutralization with 0.5 M 2-(N-morpholino)ethanesulfonic acid (pH 6.0). The proteins were added to an AAV2-immobilized 96-well plate and incubated at 30°C for 1 h. The plate was washed with TBS-T, and subsequently, horseradish peroxidase-conjugated anti-His antibody was added, followed by incubation for 1 h. After washing the plate with TBS-T, 3,3’,5,5’-tetramethylbenzidine was added to the plate to detect protein binding activity. The reaction was stopped using 1 M phosphoric acid, and chemiluminescence was measured by assessing adsorption at 450 nm.

### SPR

The interaction between WT-AAVR or AR-AAVR and AAV was analyzed using SPR with the Biacore 8K instrument (Cytiva). The proteins were dialyzed against a running buffer (0.15 M NaCl, 20 mM HEPES-NaOH, 10 mM CaCl_2_, and 0.005% Tween20; pH 7.4). A CM5 sensor chip (Cytiva) was used to immobilize AAV. AAV2 and AAV5 were immobilized on the surface of the chip at densities of 1600 and 900 RU, respectively, through amine coupling. The analyte concentrations were adjusted to 5, 2.5, 1.25, 0.625, 0.313, 0.156, and 0.078 µM. Sensorgrams corresponding to the binding of WT-AAVR or AR-AAVR to AAV were obtained by injecting increasing concentrations of the analyte at a flow rate of 30 µL/min. The contact and dissociation times were 90 s and 180 s, respectively. Regeneration was performed after the completion of each sensorgram by injecting a solution of 0.1 M glycine at pH 3.0. Data analysis was performed using the BIAevaluation Software (Cytiva). The dissociation constant (*K*_D_) was determined from a plot of steady-state binding levels against analyte concentrations.

### DSC

The thermal stabilities of the WT-AAVR and AR-AAVR were determined through DSC using a VP-DSC instrument (Malvern Panalytical, Grovewood Road, United Kingdom). Before each scan, protein samples were dialyzed using a DSC buffer (0.15 M NaCl, 20 mM Tris-HCl, and 10 mM CaCl_2_; pH 7.4). Protein samples at a concentration of 50 µM were subjected to a heating cell at 20–100°C at a scan rate of 1.0°C/min. The thermograms of the protein samples were normalized by subtracting their signal from that of the reference cell containing only the buffer. The melting temperature (*T*_m_) values were calculated by a standard fitting procedure using the Microcal Origin 7.0 software and a non-two-state model.

### Chromatography

The affinity resin in which WT-AAVR or AR-AAVR was immobilized was packed in an empty steel use stainless (SUS) column (4.6 mm × 75 mm). Each AAVR was immobilized on a non-porous methacrylate resin (5 μm) by an optimized highly orientated coupling method on residues of the immobilized tag. Another affinity column was prepared by packing the POROS™ CaptureSelect™ AAVX Affinity Resin (Thermo Fisher Scientific) into an empty SUS column (4.6 mm × 75 mm). Mobile phase A consisted of 15 mM sodium acetate, 10 mM glycine, and 50 mM CaCl_2_ (pH 4.5), while mobile phase B consisted of 15 mM sodium acetate, 10 mM glycine, and 50 mM CaCl_2_ (pH 2.2). A linear gradient of buffer B (0–100%) was applied to the column at a flow rate of 0.5 mL/min to elute AAV using an HPLC system (Shimadzu, Kyoto, Japan).

### ELISA or DLS to quantify AAV capsid titers

The AAV Titration ELISA Kit series (PROGEN Biotechnik GmbH) was used to quantify AAV capsid titers depending on the AAV serotype. To assess the AAV particle size and shape, the DLS of purified AAV was measured using the Zetasizer Ultra (Malvern Panalytical). In addition, multi-angle DLS (MADLS) was measured to quantify the capsid titer.

### AAV plasmids

The helper plasmid and the AAV-ITR plasmid with a reporter gene (pAAV-ZsGreen1) were acquired from Takara Bio. The Rep/Cap plasmids were prepared according to AAV serotypes. AAV1, AAV2, AAV5, and AAV6 were prepared using the AAVpro® Packaging Plasmid (Takara Bio, Shiga, Japan), while AAV4, AAV8, and AAV9 were synthesized by FASMAC (Kanagawa, Japan).

### AAV production and purification

AAV was produced using a Gibco AAV-MAX production system (Thermo Fisher Scientific).

The Gibco Viral Production Cells 2.0 were cultured in the Viral Production Medium supplemented with GlutaMAX (Thermo Fisher Scientific). AAV expression was induced through triple transfection, as per the AAV-MAX Transfection Kit protocol. The culture medium was centrifuged to remove the cells and obtain the cell culture supernatant, which was further filtered. The cell-lysed supernatant was obtained by adding a surfactant (Tween 20 or Triton X-100) to the culture medium and removing the debris via centrifugation and filtration.

The harvested AAV was purified via column chromatography using an affinity resin. The POROS CaptureSelect AAVX Affinity Resin (Thermo Fisher Scientific) was packed into empty Tricorn chromatography columns (Cytiva). The AAV-expressing cell-cultured or cell-lysed supernatants were applied to the column using the ÄKTA go™ chromatography instrument (Cytiva). The equilibration and washing were performed using a wash buffer (0.5 M NaCl, 10 mM CaCl_2_, and 20 mM Tris-HCl; pH 7.4), and the AAV was eluted with an elution buffer (0.5 M NaCl and 0.1 M acetate; pH 2.0). The eluted AAV was immediately neutralized with a neutralization buffer (20 mM MgCl_2_ and 1 M Tris-HCl; pH 8.5). The capsid titer of the purified AAV8 was quantified using MADLS as the control.

## Data availability

The datasets generated during the current study are available from the corresponding author upon reasonable request.

## Acknowledgments

The authors thank Ryuma Ikeura, Keita Omura, and Chiyuri Akimono of Tosoh Corporation, Life Science Research Laboratory, for their technical support.

## Author contributions

K.Y., K.K, K.W., Y.M., T.T., and T.I. conceived and designed the experiment. K.Y., K.K, K.W., and Y.M. performed the experiments. This work was supervised by Y.T, T.T., T.I., and T.O. K.Y., Y.T., and T.O. wrote the manuscript. All authors reviewed, edited, and approved the manuscript.

