## Supplemental Information for "Engineering a highly durable adeno-associated virus receptor for analytical applications"

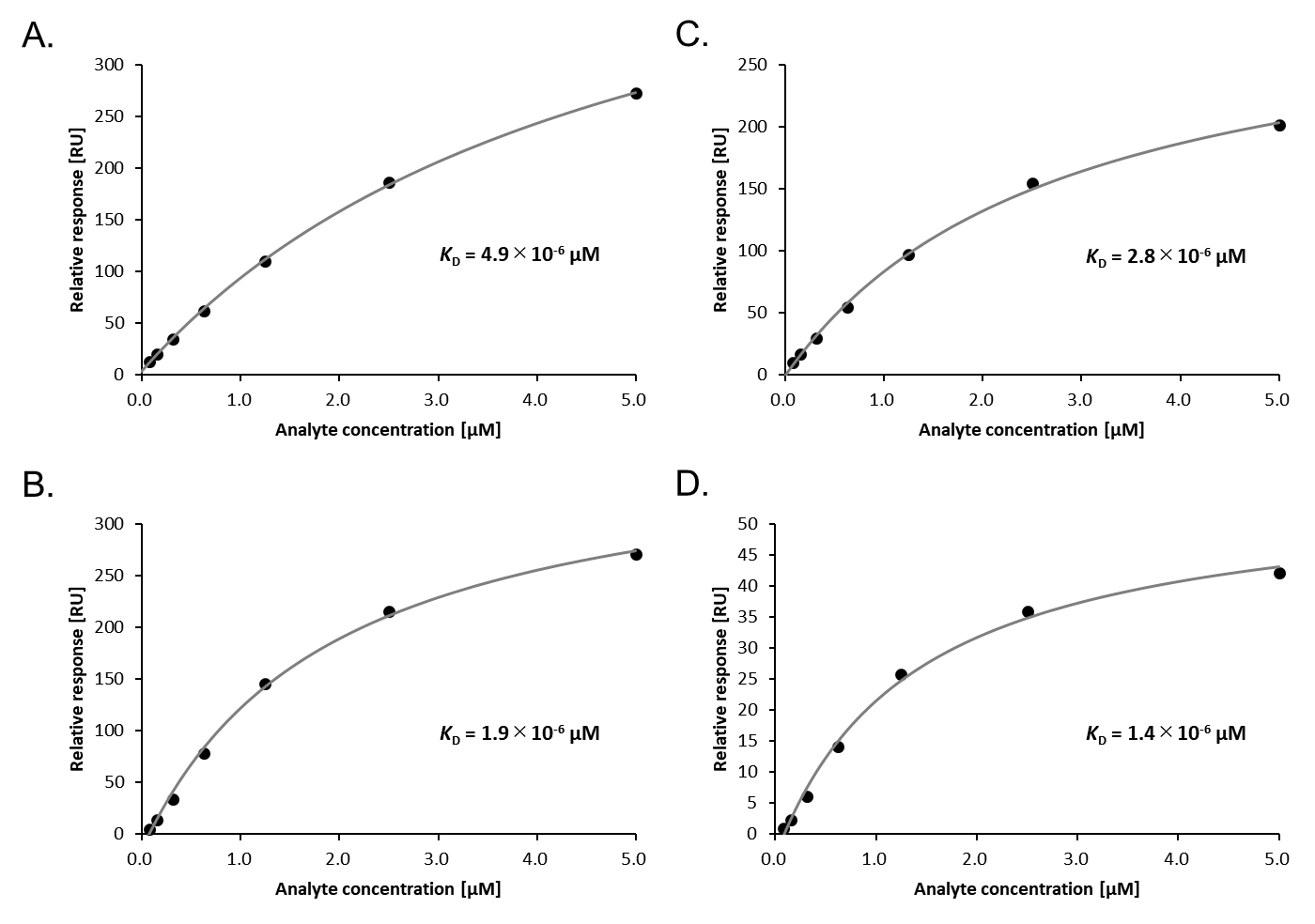

Fig. S1. Binding responses of AAVR: (A) WT-AAVR and AAV2, (B) WT-AAVR and AAV5, (C) AR-AAVR and AAV2, and (D) AR-AAVR and AAV5. The black dots and curves represent the binding response and fitted data, respectively. *K*_D_ values were obtained from the fitted curves.

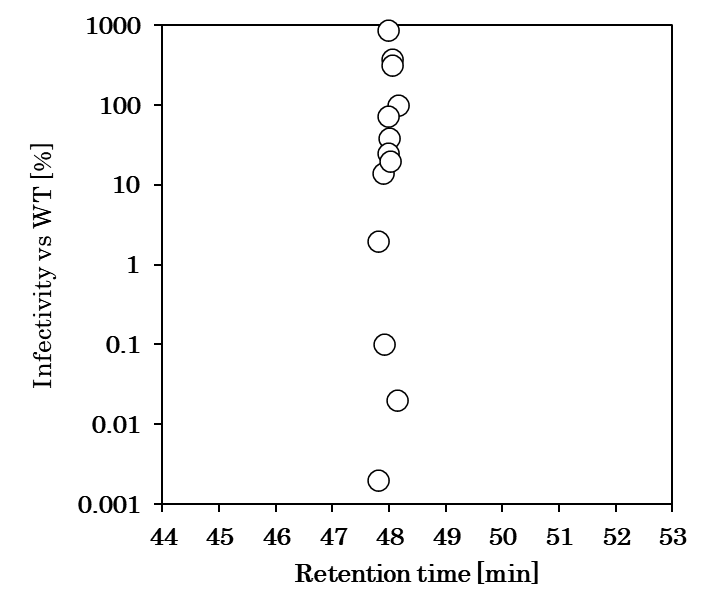

Fig. S2. Plots of retention time against the transduction efficiency of each AAV2 mutant from Table 1 when analyzed using the non-AAVR affinity column.

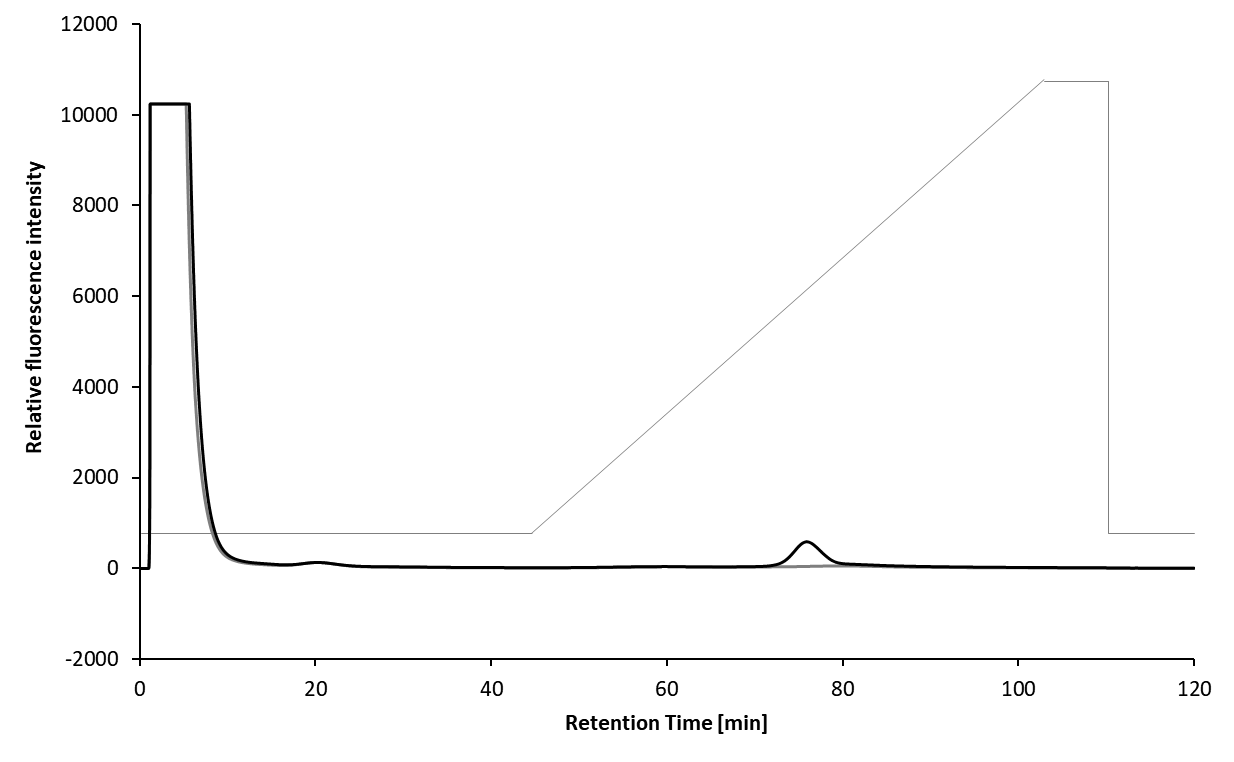

Fig. S3. Chromatograms of the cell-lysed supernatants of AAV8-expressed (black) and untreated (gray) cells analyzed using the AVR-NPR column. The gray polygonal line represents the ratio of mobile phase A (15 mM sodium acetate, 10 mM glycine, and 50 mM CaCl_2_ (pH 4.5)) and B (15 mM sodium acetate, 10 mM glycine, and 50 mM CaCl_2_ (pH 2.2)), which here the ratio of B increases linearly from 0% to 100% between 35 and 95 min after the start of the measurement.

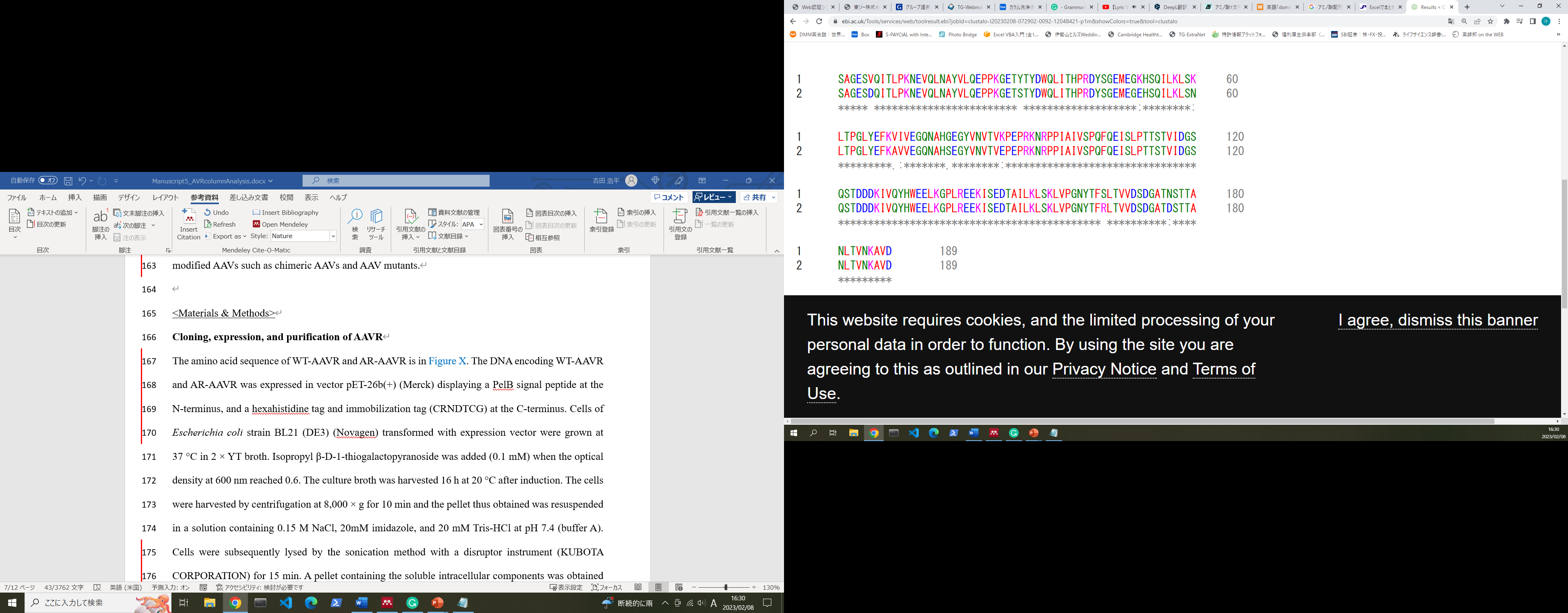

WT-AAVR

AR-AAVR

WT-AAVR

AR-AAVR

WT-AAVR

AR-AAVR

WT-AAVR

AR-AAVR

Fig. S4. Amino acid alignment of WT-AAVR and AR-AAVR generated using ClustalOmega.
